# Converging inputs compete at the lateral parabrachial nuclei to dictate the affective-motivational responses to cold pain

**DOI:** 10.1101/2024.03.25.586591

**Authors:** Prannay Reddy, Shrivas Chaterji, Austin Varughese, Yatika, Anupama Sathyamurthy, Arnab Barik

## Abstract

The neural mechanisms of the affective-motivational symptoms of chronic pain are poorly understood. In chronic pain, our innate coping mechanisms fail to provide relief leading to heightened manifestation of these behaviors. In laboratory animals, such as mice and rats, licking the affected areas is a behavioral coping mechanism and it is sensitized in chronic pain. Hence, we have focused on delineating the brain circuits mediating licking in mice with chemotherapy-induced peripheral neuropathy (CIPN). Mice with CIPN develop intense cold hypersensitivity and lick their paws upon contact with cold stimuli. We studied how the lateral parabrachial nucleus (LPBN) neurons facilitate licking behavior when mice are exposed to noxious thermal stimuli. Taking advantage of transsynaptic viral, optogenetic, and chemogenetic strategies, we observed that the LPBN neurons become hypersensitive to cold in mice with CIPN and facilitate licks. Further, we found that the expression of licks depends on competing excitatory and inhibitory inputs from the spinal cord and lateral hypothalamus (LHA), respectively. We anatomically traced the post-synaptic targets of the spinal cord and LHA in the LPBN and found that they synapse onto overlapping populations. Activation of this LPBN population was sufficient to promote licking due to cold allodynia. In sum, our data indicate that the nociceptive inputs from the spinal cord and information on brain states from the hypothalamus impinge on overlapping LPBN populations to modulate their activity and, in turn, regulate the elevated affective-motivational responses in CIPN.

## 1. Introduction

When exposed to noxious stimuli, we respond in two distinct ways: first, we reflexively withdraw or escape from the stimuli to prevent injury. Second, we attend to the wound to cope with the pain, if we do sustain an injury. Humans employ various coping strategies, including rubbing the wound, holding the affected area under running water, or seeking professional help. Similarly, laboratory mice with injured paws respond by shaking and licking^78,32,8^. While the neural mechanisms of reflexive withdrawal and escape are relatively well understood^67,97,6^, those underlying pain-directed attention and coping behaviors, such as shaking or licking, remain unclear.

In classic decerebration experiments where mid- and fore-brain structures were removed whilst preserving connections between the spinal cord and brainstem, rats could shake their paws, but not lick, in response to persistent noxious stimuli^51^. In contrast, intact rats licked and shook the affected paws in response to the same stimuli. This observation suggests that while mid- and forebrain structures are essential for expressing pain-induced licking behavior, brainstem-spinal cord circuitries are sufficient for eliciting shaking responses. Notably, decerebrated animals did not display emotional responses to pain, indicating that while both licking and shaking serve to reduce the unpleasantness of pain and can be regarded as the index of pain intensity, only licking reflects the affective-motivational or emotional component of pain sensation, while shakes appear to be a reflexive response to pain. Here, we sought to understand the central circuit mechanisms underlying pain-induced licking.

Recently, Huang et al. demonstrated the involvement of a molecularly defined neuronal population in the spinal cord that is essential for pain-induced licking behavior in mice^30,41,44,83^. These neurons express the gene encoding the neuropeptide, substance P or *Tac1,* and primarily project to the lateral parabrachial nucleus (LPBN). LPBN neurons that receive spinal inputs in turn target several areas in the mid and forebrain regions involved in emotion and motivation, including the hypothalamus, thalamus, insula, and amygdala^68,74,80^, making it a critical conduit for transmitting pain information across the brain and mediating affective responses to pain, including licking behavior^12,81,89,22,5^. Indeed, activating LPBN neurons with strong projections to the intralaminar thalamus promotes licking in response to sustained noxious mechanical and chemical stimuli^5,27^. On the other hand, LPBN neurons receive reciprocal inputs from the brain regions such as the amygdala and hypothalamus^70,7,55,91,60^. Many of these inputs are inhibitory and are instrumental in facilitating the pain-suppression effects of physiological brain states such as hunger and stress^71,50,39,38^. For example, Agrp neurons housed in the arcuate nucleus are activated by hunger and in turn, inhibit LPBN nociceptive neurons^19,93,34^. Similarly, amygdala neurons that respond to anesthetics may impart their analgesic effects by inhibiting neurons in the PBN^42^. Thus, LPBN-mediated expression of pain-induced licking behaviors is driven by nociceptive information from the dorsal horn of the spinal cord and is simultaneously gated by the descending inputs from the fore/midbrain regions that encode internal brain states. Mice do not lick their paws spontaneously but do so in response to sustained noxious stimuli^66,90^. In particular, laboratory techniques that are used to induce acute and chronic peripheral sensitization, such as intraplantar injections of AITC, CFA, capsaicin, or formalin, or exposure to innocuous or noxious mechanical or thermal stimuli induce vigorous licking^1,17,75,84^. Here, we investigated the circuit mechanisms underlying licking responses elicited by cooling stimuli in mice with chemotherapy-induced peripheral neuropathy (CIPN), a common and challenging side-effect of cancer chemotherapy treatment in humans^20,37,57,58^. Upon intra-plantar injection (i.pl.) of oxaliplatin, mice develop cold hypersensitivity and exhibit vigorous licks^28,53^ in response to cooling stimuli. We investigated how the LPBN may play a role in cold-induced defensive behaviors in mice with CIPN and found that the LPBN facilitates licking in response to innocuous cold stimuli in mice with CIPN. In addition to receiving ascending somatosensory inputs from the spinal cord, LPBN neurons also receive descending inputs from neurons in the lateral hypothalamus (LHA), which are known to encode information on the internal states of the brain, such as hunger and stress^4,13,34,43,46,85^, and these LHA inputs to the LPBN suppress the licking behaviors^2^. In summary, how LPBN neurons shape the expression of coping behaviors such as licking in response to cold in mice with neuropathy is decided by the competing inputs from the spinal cord and the LHA.

## 2. Methods

### 2.1. Mouse lines

Animal care and experimental procedures were performed following protocols approved by the CPCSEA at the Indian Institute of Science. The animals were housed at the IISc Central Animal Facility under standard animal housing conditions: 12 h light/dark cycle from 7:00 am to 7:00 pm with ad-libitum access to food and water; mice were housed in IVC cages in Specific pathogen-free (SPF) clean air rooms. Mice strains used: Vglut2-Cre or Slc17a6tm2(Cre) Lowl/J(Stock number 016963); Ai14 (B6;129S6-Gt(ROSA)26Sortm9(CAG-tdTomato)Hze/J (Stock No 007905), BALB/cJ, Jackson Laboratories, USA. Experimental animals were between 2-4 months old.

### 2.2. Viral vectors and stereotaxic injections

Mice were anesthetized with 2% isoflurane/oxygen before and during the surgery. The craniotomy was performed using a handheld micro drill. The viral injections were administered using a hamilton syringe (10 ul) with glass-pulled needles. The viral injections (1:1 in saline) were 300 nL, infused at a rate of 100 nL/min. The following were the coordinates for viral injections: Pbn (AP: −5.30 ML: +1.50 DV: −3.15); LHA (AP: −1.70 ML: +1.00 DV: −5.15). The stereotaxic surgeries to deliver AAVs in the lumbar spinal cord were performed as described previously in [10]. The vectors used and their sources: AAV1-hSyn-Cre.WPRE.hGH (Addgene, Catalog# v126225); AAVretro-pmSyn1-EBFP-Cre (Donated by Ariel Levine, NIH); scAAV-1/2-hSyn1-FLPO-SV40p(A) (University of Zurich, Catalog# v59-1); ssAAV-9/2-hGAD67-chl-icre-SV40p(A) (University of Zurich, Catalog# v197-9); pAAV5-hsyn-DIO-EGFP (Addgene, Catalog# 50457-AAV 1); pAAV5-FLEX-tdTomato (Addgene, Catalog# 28306-PHP.S); pENN.AAV5.hSyn.TurboRFP.WPRE.RBG (Addgene, Catalog# 10552-AAV1); pAAV5-hsyn-DIO-hM4D(GI)-mCherry (Addgene, Catalog# 44362-AAV5); AAV9.syn.flex.GcaMP6s (Addgene, Catalog# pNM V3872TI-R(7.5)); AAV5-hSyn-hM3D(Gq)-mCherry (Addgene, catalog#50474); and pAAV-EF1a-double floxed-hChR2(H134R)-GFP-WPRE-HGHpA(Addgene, Catalog# v64219). For rabies tracing experiments, rAAV5-EF1α-DIO-oRVG (BrainVTA, Catalog# PT-0023) and rAAV5-EF1α-DIO-EGFP-T2A-TVA (BrainVTA, Catalog# PT-0062) were injected first, followed by RV-EnvA-Delta G-dsRed (BrainVTA, Catalog# R01002) after 2 weeks. Tissue was harvested after 1 week of rabies injection for histochemical analysis. Post-hoc histological examination of each injected mouse was used to confirm that viral-mediated expression was restricted to target nuclei.

Spinal cord stereotaxic injections were performed as previously reported in^21^. The tissue was harvested 1 week after the rabies virus injection. Post hoc histological examination of each injected mouse was used to confirm the nuclei targeted viral expression.

### 2.3. Fiber Photometry

A two-channel fiber photometry system from RWD was used to collect data. Two light LEDs (410/470nm) were passed through a fiber optic cable coupled to the cannula implanted in the mouse. Fluorescence emission was acquired through the same fiber optic cable onto a CMOS camera. The photometry data was analyzed using the RWD photometry software and csv files were generated. The start and end of stimuli were timestamped. All trace graphs were plotted from CSV files using GraphPad Prism 7/9 software. All heatmaps were plotted from CSV files using custom Python scripts.

### 2.4. Fiber implantation

For fiber photometry and optogenetics, fiber optic cannulas from RWD with Ø1.25 mm Ceramic Ferrule, 200 μm Core, 0.22 NA, and L = 5 mm were implanted at the following coordinates after infusion of the: LPBN (AP:-5.30 ML:+-1.50 DV:-3.15); LHA (AP:-1.70 ML+-1.00 DV:-5.15). The fibers were implanted after infusion of either GCaMP or Chr2 in LPBN or LH. Animals were allowed to recover for at least 3 weeks before performing behavioral tests. Successful labeling and fiber implantation were confirmed post hoc by staining for GFP/mCherry for viral expression and injury caused by the fiber for implantation. Animals with viral-mediated gene expression at the intended locations and fiber implantations, as observed in post hoc tests, were only included.

### 2.5. Behavioral assays

Behavioral assays for the same cohorts were handled by a single experimenter. Before experimentation, the experimenter was blinded to the identity of animals. Mice were habituated in their home cages for at least 30 minutes in the behavior room before experiments. An equal male-to-female ratio was maintained in every experimental cohort and condition unless otherwise stated, with no significant differences seen between sexes in the responses recorded from the behavioral experiments. Wherever possible, efforts were made to keep the usage of animals to a minimum. Oxaliplatin (2mg/ml) was loaded into an insulin syringe, 20µl was injected into the left paw, and the mice were left in their home cages in the behavior room for a minimum of 4 hours before any behavioral experiments to cause persistent inflammatory pain and thermal hypersensitivity. For chemogenetic experiments, Clozapine-N-Oxide (1mg/kg) dissolved in DMSO was diluted in saline and injected intraperitoneally at least 20-30 minutes before behavioral experiments. For optogenetic experiments, the fiber-coupled LED (channelrhodopsin activation; RWD, China) was connected to the optic ferrules implanted on the heads of mice. The mice were left for 20 minutes to habituate to the fiber and were then placed in their respective chamber depending on the behavioral assay. During experiments, the mice received photo stimulation with a power of 20mW at 15Hz with a pulse width of 5ms. For the cold plate, hot plate, and gradient hot plate assay, the mice were placed on the cold plate covered by an acrylic box. The hot-cold analgesia plate was purchased from Orchid Scientific and was used according to the manufacturer’s instructions. The cold plate assay lasts for 5 minutes, where the starting temperature is 24 degrees, and it drops at 4.8 degrees/minute until it reaches 0 degrees. The hotplate assay lasts for 45 seconds, where the temperature is set at 52 degrees. The gradient hot plate assay lasts for 5 minutes, where the starting temperature is 32 degrees and it increases at 4.0 degrees/minute until it reaches 52 degrees. For the cold air puff assay, the mice were placed on an elevated metal mesh surface and were covered in an acrylic box to restrict their movement. Dry ice was finely crushed and loaded into a 20ml syringe, after which air was puffed from below towards the hind paws of mice. A minimum of 10 trials for each paw of the mice were conducted. All the experiments were videotaped simultaneously with three wired cameras (Logitech, USA) placed horizontally and scored offline post hoc manually. All experiments were scored by an experimenter blinded to the different experimental conditions.

### 2.6. Immunostaining

Mice were anesthetized with isoflurane and perfused intracardially with 1X Phosphate Buffered Saline (PBS) (Takara, Japan) and 4% Paraformaldehyde (PFA) (Ted Pella, Inc., USA), consecutively for immunostaining experiments. For hM3Dq cFos experiments, brains and spinal cords were harvested 150 minutes after i.p. DCZ administration. For hM4Di cFos experiments, 60 minutes after i.p. DCZ administration, the mice were placed on the cold plate, 90 minutes after which their brains and spinal cords were harvested. The brains and spinal cords were harvested and further fixed in 4% PFA overnight at 4°, then stored in 30% sucrose overnight. The brains and cords were embedded in OCT medium and were cut at a thickness of 50um sections using a cryostat. Spinal sections were directly mounted onto glass slides, whereas brain sections were floated in PBS and mounted onto glass slides before staining. Tissue sections were rinsed in 1X PBS and incubated in a blocking buffer (2% Bovine Serum Albumin (BSA); 0.3% Triton X-100; PBS) for 1 hour at room temperature. Sections were incubated in primary antibodies in a blocking buffer at room temperature overnight. Sections were rinsed 1-2 times with PBS and incubated for 2 hours in Alexa Fluor conjugated goat anti-rabbit/ chicken or donkey anti-goat/rabbit secondary antibodies (Invitrogen) at room temperature, washed in PBS, and mounted in VectaMount permanent mounting media (Vector Laboratories Inc.) onto charged glass slides (Globe Scientific Inc.). We used an upright fluorescence microscope (Khush, Bengaluru) (2.5X, 4X, and 10X lenses) and ImageJ/FIJI image processing software to image, and process images for the verification of anatomical location of cannulae implants. For the anatomical studies, the images were collected with 10X and 20X objective on a laser scanning confocal system (Leica SP8 Falcon, Germany) and processed using the Leica image analysis suite.

### 2.7. Statistical analysis

All statistical analyses were performed using GraphPad Prism 7/9 software. ns>0.05, *<0.05, **<0.01, ***<0.001, ****<0.0001.

### 2.8. Data availability

Raw data will be made available upon request.

## 3. Results

### 3.1. Intraplantar Oxaliplatin induces long-term cold allodynia

To delineate the central circuits for affective-motivational behaviors in mice, particularly paw-licking responses to sustained pain, we initially induced cold allodynia in mice through intraplantar administration of oxaliplatin(Fig. 1A), a commonly used chemotherapy drug that is known to induce chemotherapy-induced peripheral neuropathy (CIPN) along with cold and mechanical hypersensitivity ^11,35,48,64^. Oxaliplatin administration resulted in increased licks and shakes on the gradient cold plate test, where the surface temperature was gradually reduced from 24°C to 0°C (Fig.1B). Notably, the licks and shakes appeared as the temperature dropped below 20°C — a range typically considered innocuous for mice. In contrast, mice injected with i.pl. saline did not shake or lick their paws on the gradient cold plate test (Fig. 1B). Conversely, in the hot-plate test set at 52°C the number of licks and shakes was comparable between the saline and oxaliplatin-injected animals (Fig. 1C). Thus, i.pl. oxaliplatin sensitized mice to cold stimuli leading to vigorous shaking and licking of the injected paw in response to innocuous cooling stimuli. Interestingly, the number of licks increased proportionally with a fall in temperature of the cold plate (Fig. 1D). However, shakes did not follow a similar trend (Fig. 1E). When expressed as a ratio of licks to shakes, the emotional response proportionally increased with decreasing cold-plate temperatures (Fig. 1F). Thus, the emotional coping response to cold pain indicated the intensity of pain in the form of increased licks. Cold hypersensitivity persisted for a week post-oxaliplatin injection, indicating sustained cold allodynia (Fig. 1G & H); however, both licks and shakes were evoked in similar proportions (Fig. 1I) during this time. Thus, the cold hypersensitivity mediated nocifensive behaviors of both reflexive and emotional nature are sustained proportionally for days after the induction of neuropathicpain.

**Fig 1. Intra-plantar.**
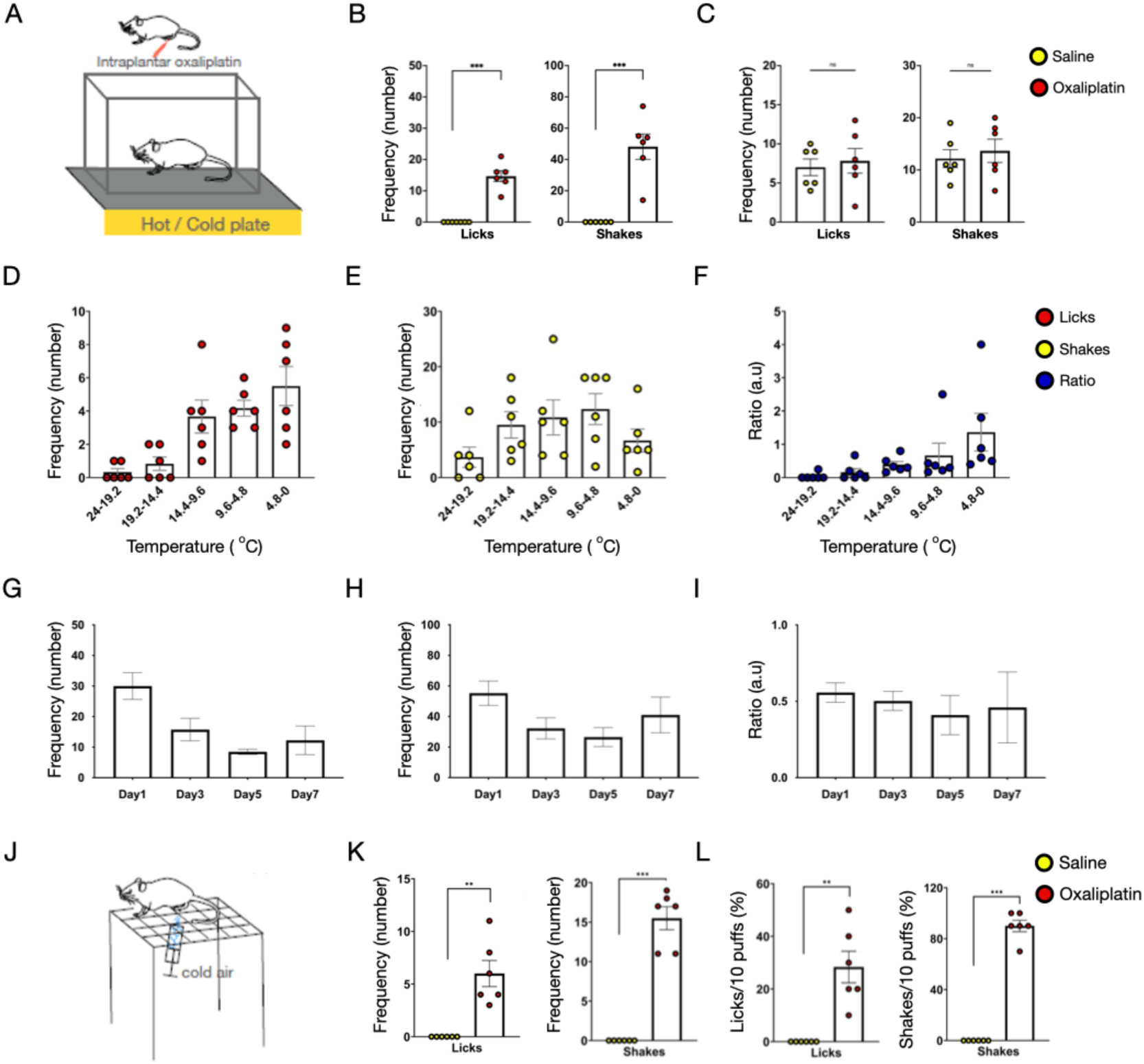
injection of oxaliplatin induces cold allodynia. (A) A schematic of cold/hot plate assay post-CIPN induction in the left hind paw of mice by intraplantar (i.pl.) oxaliplatin injection (20µl of 5 mg/ml). (B) Mice with i.pl. oxaliplatin responded with licks (14.67 ± 1.75) (\*\*\**p* = 0.0004) (left) and shakes (48.17 ± 8.11) (\*\*\**p* = 0.0019) (right) on the cold plate assay. No licks or shakes were seen in mice with i.pl. Saline. (t-test \*\*\**p* < 0.001, n=6) (C) Mice with i.pl. saline demonstrated comparable licks (7.00 ± 1.07) (left) and shakes (12.17 ± 1.72) (right) on the hot plate assay compared to the licks (7.83 ± 1.58) (left) and shakes (13.67 ± 2.23) (right) of mice with i.pl. Oxaliplatin (n=6). (D) In mice with i.pl. Oxaliplatin, the number of licks per minute on the cold plate assay were plotted against the drop in temperature on a gradient cold-plate test and the number of licks were as follows: 0.33 ± 0.21 licks in the first minute with a temperature drop from 24 to 19.2°, 0.83 ± 0.40 licks in the second minute with a temperature drop from 19.2 to 14.4°, 3.67 ± 0.99 licks in the third minute with a temperature drop from 14.4 to 9.6°, 4.17 ± 0.48 licks in the fourth minute with a temperature drop from 9.6 to 4.8°, and 5.50 ± 1.18 licks in the fifth minute with a temperature drop from 4.8 to 0° (n=6). (E) The number of shakes per minute on the cold plate assay with the same drop (D) in temperature were as follows: 3.67 ± 1.82 shakes in the first minute, 9.50 ± 2.38 shakes in the second minute, 10.83 ± 3.15 shakes in the third minute, 12.33 ± 2.79 shakes in the fourth minute, and 6.68 ± 2.08 shakes in the fifth minute (n=6). (F) The lick-to-shake (affective-motivational:reflexive coping responses) ratios per minute on the cold plate assay with the same drop in temperature were as follows: 0.04 ± 0.04 in the first minute, 0.16 ± 0.1 in the second minute, 0.39 ± 0.09 in the third minute, 0.66 ± 0.37 in the fourth minute, and 1.36 ± 0.57 in the fifth minute (n=6). (G, H) Innocuous cold induced licks and shakes lasted for more than a week post-ipl. Oxaliplatin administration: Day 0 as the day of injection were as follows: 30.00± 4.39 on Day 1, 15.75± 3.68 on Day 3, 8.50± 0.87 on Day 5, and 12.25± 4.70 on Day 7 (n=4). The number of shakes (H) over the week on the cold plate assay with Day 0 as the day of injection were as follows: 55.25 ± 7.9 on Day 1, 32.25 ± 6.91 on Day 3, 26.5 ± 6.20 on Day 5, and 41.0 ± 11.67 on Day 7 (n=4). (I) The lick-to-shake ratios throughout the week on the cold plate assay with Day 0 as the day of injection were as follows: 0.56 ± 0.06 on Day 1, 0.50 ± 0.06 on Day 3, 0.41 ± 0.13 on Day 5, and 0.46 ± 0.23 on Day 7 (n=4). (J) A schematic of the cold air puff assay. (K) Comparison of the lick responses (left) after i.pl. saline (0.00 ± 0.00) and oxaliplatin (6.00 ± 1.24) (\*\**p* = 0.0047); and shakes (right) after i.pl. saline (0.00 ± 0.00) and oxaliplatin (15.50 ± 1.46) (\*\*\**p* = 0.0001) during 10 trials of the cold air puff assay trials including spontaneous licks and shakes between trials. (n=6). (L) Comparing the percentage of licks (left) after i.pl. saline (0.00 ± 0.00%) and oxaliplatin (28.3 ± 6.0%) (\*\**p* = 0.0053); and shakes (right) after i.pl. saline (0.00 ± 0.00%) and oxaliplatin (90.0 ± 4.5%) (\*\*\**p* = 0.0001) during 10 trials of the cold air puff assay. (t-test \*\*\**p* < 0.001, and \*\**p* < 0.01, n=6).

Further, we probed if transient cooling stimuli, such as cold air puffs directed towards the paw with neuropathic pain, could generate nocifensive behaviors such as licks and shakes. Unlike the cold plate test, where the cooling stimulus persisted even after the mice reacted, in the cold puff test, stimulus application was terminated upon the onset of nocifensive behaviors. A similar application of transient mechanical stimuli with von Frey filaments or thermal stimuli with the Hargreaves apparatus induces rapid withdrawal, shakes, and licks in mice with peripheral neuropathy or inflammation^9,82,59,96^. When cold air between ∼6°C and 10°C (Fig. S1A), which is in the innocuous range, was applied to the paws with neuropathic pain, mice responded with licks and shakes (Fig. 1K & L). The frequency of licks and shakes due to the cold air puff was comparable, indicating that the transient cold stimulus was sufficient to mediate reflexive and emotional coping responses and suggesting that (Fig. S1B). This suggests that a transient cold stimulus could recruit and sustain activity in the supraspinal circuits to mediate coping behaviors.

Next, we tested if a single cold puff that can generate nocifensive responses to cold pain could engage the spino-parabrachial pathway. As predicted, application of cold air puff on the paws of mice with CPIN resulted in increased expression of c-Fos, an immediate early gene (IEG), both in neurons in the superficial dorsal horn (Fig. S1C), which is the chief source of spinal inputs to the lateral parabrachial nucleus (LPBN) [87][92] and the LPBN (Fig. S1D).. In contrast, there were fewer c-fos expressing neurons in the spinal cord or the LPBN of saline injected mice. Overall, these results suggested that transient innocuous cold stimuli can evoke reflexive and affective-motivational behavioral responses in mice with CIPN and potentially involve the spino-parabrachial pathway.

### 3.2. LPBN neurons are engaged by cold allodynia

As noted previously, the spino-parabrachial pathway is engaged by innocuous cold stimuli in mice with CIPN (Fig. S1C, D). However, monitoring neural activity solely through cFos expression provides a static view of the stimuli -evoked neural ensembles in the LPBN. To gain insight into LPBN dynamics in awake-behaving mice experiencing cold pain, we employed fiber photometry^24^, a technique, which allows optical recording of neural activity from deep brain regions expressing genetically encoded calcium sensors such as GCaMP6s^25,49^. Since most LPBN neurons are glutamatergic^6,40,68^, we selectively expressed GCaMP6s in the LPBN of VGlut2-Cre mice (reference) using stereotaxically delivered AAVs^25,49^ (Fig. 2A to C). We recorded from the ipsi- and contra-lateral (to the oxaliplatin or saline-injected paw) LPBN, since both receive spinal inputs from one side of the spinal cord^15,27^. As expected, we observed increases in calcium transients in both the ipsi- and contra-LPBN neurons with decreasing temperatures on the cold plate test and with rising temperatures on the gradient hot-plate test(Fig. 2D to F)(Fig.S2) in neuropathic but not control mice. Noxious heat activates LPBN neurons and elicits nocifensive behaviors such as licks and shakes^33,86,29^. Notably, while innocuous cold engaged LPBN neurons in mice with CIPN, innocuous heat did not (Fig. 2D), implying that the mice were sensitive specifically to cold and not heat stimuli. Further, we correlated LPBN activity with licks and shakes on the gradient cold-plate and hot-plate tests and found similar magnitudes of LPBN activation during both these responses (Fig. 2G to P), indicating that the LPBN neurons were engaged in both spinal and supraspinal coping responses to cold pain. Further analysis showed an increase in the occurrence of high amplitude calcium transients (>3 times the Median Absolute Deviation) in the LPBN of CIPN mice in the cold-plate test as compared to controls, and a decrease in the occurrence of high amplitude calcium transients in the hot-plate test (Fig. 2Q & R). This observation implies that the CIPN may render the LPBN neurons more sensitive to cold than heat in the noxious range. Next, we tested if the LPBN neurons in neuropathic mice are activated by cold air puff. The air-puff test allowed us to direct the cold stimuli to the injured paw and provide a transient stimulus (Fig. 1J). As observed in the cold-plate test, we found that cold air puff on the injured paw in the neuropathic mice consistently engaged both ipsi- and contra-lateral LPBN neurons (Fig. 2S-X). Interestingly, the activity achieved maximum amplitude post-termination of the cold air stimuli (Fig. 2T, W), in agreement with the behavior observed in the cold-puff test (Fig. 1K & L). Our fiber photometry recordings suggest that the LPBN neurons are sensitized to peripheral stimuli in mice with CIPN and play an essential role in mediating nocifensive responses to cold pain.

**Fig 2.**
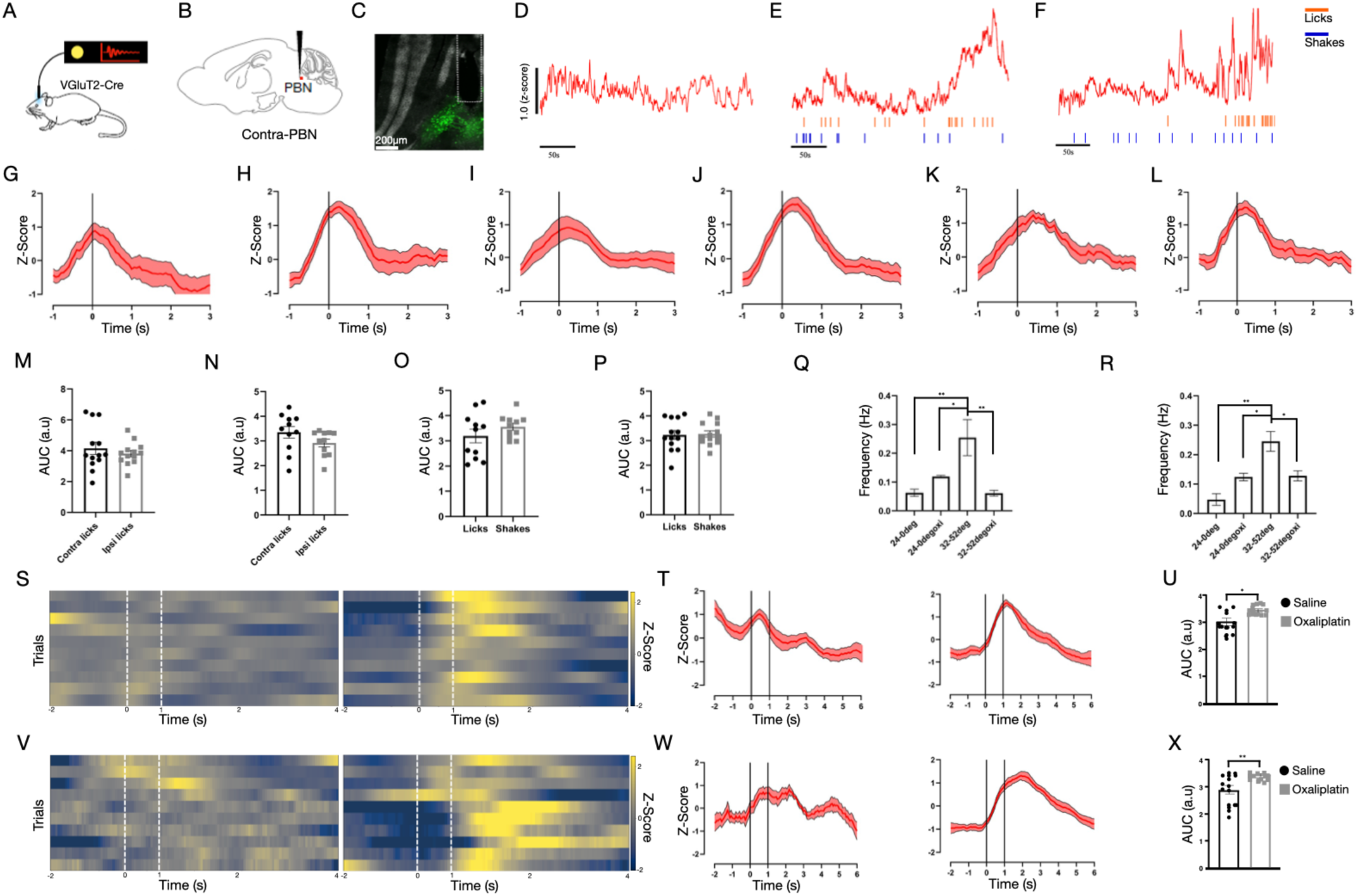
LPBN neurons are engaged during cold allodynia. (A) A schematic of the fiber photometry assay used to record neural activity from the LPBNof VGlut2-Cre transgenic mouse line. (n=3 mice for both sides.) (B) A schematic of a viral mediated Cre-dependent expression of the genetically encoded calcium indicator GCaMP6s in the LPBN of VGlut2-cre transgenic mice with a fiber optic cannulae implanted over the LPBN. (C) A representative confocal image of the expression of GCaMP6s (green) with thefiber placement (white dotted line). (D) A representative trace of the calcium activity in the LPBN^VGlut2^ neurons during the gradient cold plate assay after i.pl. saline, (E) i.pl. Oxaliplatin, (F) and in the gradient hot plate assay where the temperature increases from 32° to 52° at the rate of 4° C/min post i.pl. oxaliplatin. The orange and blue lines below are incidences of licks and shakes, respectively. (G) The average traces of calcium responses based on normalized Z-score values recorded from the LPBN^VGlut^^2^ neurons during gradient hotplate assay after i.pl. saline for ipsilateral hind paw licks (H) and contralateral hind paw licks (t = 0s is the onset of the response) (trials=15, n=3). (I) The average traces of calcium responses based on normalized Z-score values recorded from the LPBN^VGlut2^ neurons during cold plate assay after i.pl. oxaliplatin during ipsilateral hind paw licks (J) and ipsilateral hind paw shakes, (K) contralateral hind paw licks, and (L) contralateral hind paw shakes (t = 0s is the onset of the response) (trials=15, n=3). (M) A comparison of the neural activity of LPBN^VGlut2^ neurons between contralateral and ipsilateral licks in gradient hotplate after i.pl. saline with mean area of 4.15 ± 0.40 and 3.81 ± 0.21 respectively and (N) between contralateral and ipsilateral licks in gradient hotplate after i.pl. oxaliplatin with a mean area of 3.35 ± 0.24 and 2.92 ± 0.16 respectively. (trials=13, n=3). (O) A comparison of the LPBN^VGlut2^ neuronal activity between ipsilateral licks (3.19 ± 0.27) and shakes (3.56 ± 0.14); (P) between contralateral licks (3.23 ± 0.18) and shakes (3.27 ± 0.13) in cold plate assay after i.pl. oxaliplatin. (trials=13, n=3). (Q) The frequency of peaks of the Z-Score of the LPBN^VGlut2^ activity for the first 2 minutes of the recordings compared between gradient cold-plate test, with i.pl. saline (0.06 ± 0.01) (\*\**p* = 0.0095) and i.pl. oxaliplatin (0.12 ± 0.00) (\**p* = 0.035), and gradient hot-plate test with i.pl. saline (0.25 ± 0.08) and i.pl. oxaliplatin (0.06 ± 0.01) (\*\**p* = 0.0059); (R) for the last 3 minutes of recording at 14° to 0° for gradient cold plate, with i.pl. saline (0.05 ± 0.02) (\*\**p* = 0.0032) and i.pl. oxaliplatin (0.12 ± 0.01) (\**p* = 0.030), and gradient hotplate, with i.pl. saline (0.25 ± 0.03) and i.pl. oxaliplatin (0.13 ± 0.02) (\**p* = 0.026). The threshold for peaks is 3 times the median absolute deviation (MAD) value of the Z-Score of calcium activity. (One-way ANOVA \*\**p* < 0.01, and \**p* < 0.05, n=3). (S) The ipsilateral LPBN^VGlut2^ activity represented as heatmaps during cold air-puff assay for trials after i.pl. saline and oxaliplatin with normalized Z-score values (the white dotted lines indicate the duration of the cold air puff which lasted for 1 second). (trials=10, n=3). (T) The average traces of calcium responses based on normalized Z-score values recorded from the ipsilateral LPBN^VGlut2^ neurons during cold air puff assay after i.pl. saline (left) and oxaliplatin (right) (t = 0s is the onset of the cold air puff which lasts for 1s on average). (trials=15, n=3). (U) A comparison of the neural activity of ipsilateral LPBN^VGlut2^ neurons during cold air puff trials between i.pl. saline (3.04 ± 0.14) and oxaliplatin (3.44 ± 0.06) (\**p* = 0.016) (t-test \**p* < 0.05, trails=15, n=3). (V) The representative trials for the heatmaps of cold air puff assay for contralateral LPBN^VGlut2^ neuronal activity for trials after i.pl. saline and oxaliplatin with normalized Z-score (the white dotted lines indicate the duration of the cold air puff which lasts for 1s). (trials=10, n=3). (W) The average traces of calcium responses based on normalized Z-score values recorded from the contralateral LPBN^VGlut2^ neurons during cold air puff assay after i.pl. saline (left) and oxaliplatin (right) (t = 0s is the onset of the cold air puff). (trials=15, n=3). (X) A comparison of the neural activity of contralateral LPBN^VGlut2^ neurons during cold air puff trials between i.pl. saline (2.92 ± 0.16) and oxaliplatin (3.38 ± 0.03) (\*\**p* = 0.0091). (t-test \*\**p* < 0.01, trials=15, n=3).

### 3.3. LPBN neurons promote protective responses to innocuous cold in mice with cold allodynia

LPBN neurons are sufficient to promote defensive responses to noxious stimuli, for example, jumping or evading responses to noxious heat and scratching to itch^6,61^. We hypothesized that the transient activation of the LPBN neurons would increase the frequency of licks and shakes in response to cooling stimuli in mice with CIPN. To activate LPBN neurons, we expressed the gene encoding the designer receptor exclusively activated by designer drugs (DREADDs)^72^, hM3Dq, in the left LPBN (LPBN^hM3Dq^) of wild-type mice (Fig. 3A). Intraperitoneal (i.p.) administration deschloroclozapine (DCZ)^62^ in LPBN^hM3Dq^ mice increased the expression of cFos in the LPBN (Fig. 3B, Right), confirming successful activation. A similar increase in cFos expression was not observed in control LPBN^hM3Dq^ mice administered with saline (Fig. 3B, Left). We induced CIPN by injecting oxaliplatin in the right paw of LPBN^hM3Dq^ mice and observed an increase in both licking and shaking of the injected paw in DCZ injected mice compared to saline-injected controls in the gradient cold-plate (24-0°C in 300 seconds) (Fig. 3C & D), cold puff (Fig. 3E)., and hot-plate (52°C for 45 seconds) (Fig. 3F) tests. Thus, contralateral LPBN activation was sufficient to drive both reflexive and emotional coping responses to cold and hot pain. Similarly, LPBN stimulation in mice without CIPN increased licks and shakes on the hot-plate test. However, the proportion of licks and shakes were altered upon LPBN activation only on the hot-plate but not the cold plate test, suggesting that LPBN can regulate distinct coping responses depending on the nature of the stimuli. Overall, our results demonstrate that LPBN neurons play a facilitatory role in the expression of licks and shakes in response to innocuous cold in CIPN mice.

**Fig 3.**
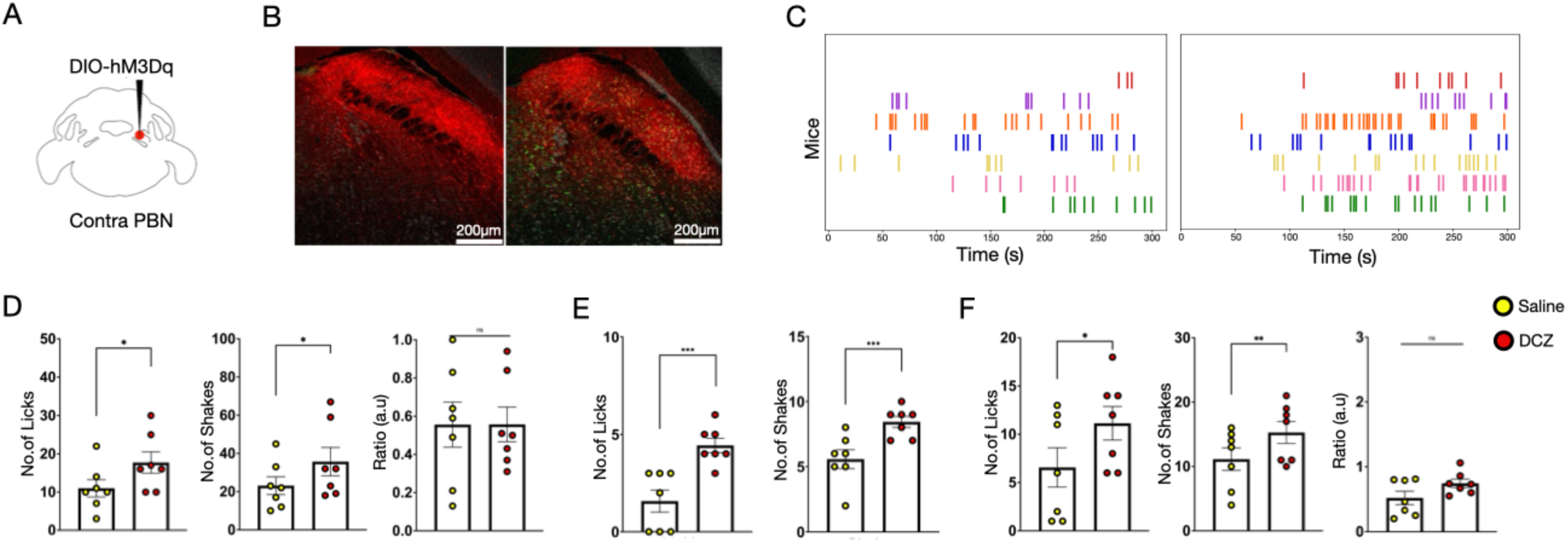
Stimulation of the LPBN promotes both licking and shake responses. (A) A schematic of DIO-hM3Dq injection in the LPBN. (n=7). (B) Representative confocal images of the expression of hM3Dq (red) and cFos (green)in LPBN after intraperitoneal (i.p.) saline (left) and i.p. DCZ injection (right). (C) Raster plots depicting the licks during cold plate assay where each row is from an individual mouse after i.p. saline (left) and after i.p DCZ (right). (D) Chemogenetic activation of the LPBN after i.pl. oxaliplatin during the cold plate assay. Comparing the licks (left) after i.p. saline (10.86 ± 2.25) and i.p. DCZ (17.71 ± 2.80) (\**p* = 0.019); the shakes (middle) after i.p. saline (23.14 ± 4.66) and i.p. DCZ (35.71 ± 7.41) (\**p* = 0.021); and the lick-to-shake ratio (right) after i.p. saline (0.56 ± 0.12) and i.p. DCZ (0.56 ± 0.09). (t-test \**p* < 0.05, n=7). (E) Chemogenetic activation of the whole LPBN after i.pl. oxaliplatin during the cold air puff assay. Comparing the licks (left) after i.p. saline (1.57 ± 0.58) and i.p. DCZ (4.43 ± 0.37) (\*\*\**p* = 0.0002); and the shakes (middle) after i.p. saline (5.57 ± 0.72) and i.p. DCZ (8.43 ± 0.43) (\*\*\**p* = 0.0004). (t-test \*\*\**p* < 0.001, n=7). (F) Chemogenetic activation of the whole LPBN after i.pl. oxaliplatin during the hot plate assay. Comparing the licks (left) after i.p. saline (6.57 ± 2.03) and i.p. DCZ (11.14 ± 1.74) (\**p* = 0.045); the shakes (middle) after i.p. saline (11.14 ± 1.75) and i.p. DCZ (15.29 ± 1.70) (\*\**p* = 0.006); and the lick-to-shake ratio (right) after i.p. saline (0.52 ± 0.10) and i.p. DCZ (0.74 ± 0.07). (t-test \*\**p* < 0.01, and \*\**p* < 0.05, n=7).

### 3.4. LPBN neurons receive excitatory inputs from the spinal cord and inhibitory inputs from the lateral hypothalamus

In addition to the excitatory projections from the spinal cord, the LPBN also receives inhibitory inputs from hypothalamic areas that can modulate pain thresholds^94,93^. To identify the specific hypothalamic areas that project to the LPBN, we injected AAVRetro-Cre in the LPBN of LSL-tdTomato transgenic mice (Fig. 4B). As expected, we found tdTomato +ve cell bodies in the LHA (Fig. 4C), in agreement with previous reports. LPBN is known to receive both excitatory and inhibitory projections from the LHA^79^. We focused on the inhibitory LHA neurons so as to define the roles of descending inhibitory inputs to PBN in the expression of nocifensive behaviors, as they may counteract the excitatory spinal inputs. To label the inhibitory neurons in the LHA, we co-injected AAV-Gad67-Cre and AAV-DIO-GFP in the LHA (Fig. 4E). As predicted, LHA Gad67 neurons innervated the lateral and medial parts of the PBN (Fig 4F). Next, we determined if the LHA inhibitory and the spinal excitatory inputs terminate in similar areas of the LPBN. To that end, we injected AAVs encoding turboRFP in the dorsal of the horn of the lumbar spinal cord and GFP driven by the Gad67 promoter^95^ in the LHA of the same mice. LHA axon terminals were spread across the entire dorsoventral and rostrocaudal axis of the LPBN, while the lumbar axon terminals were restricted in the superior-lateral and dorsolateral domains (Fig. 4H). Interestingly, in the LPBN area where spinal neurons terminate, the density of the Gad67+ve LHA axon terminals was relatively lower (Fig. 4H), suggesting that the termination zone of lumbar and LHA inhibitory projects partially overlap.

**Fig 4.**
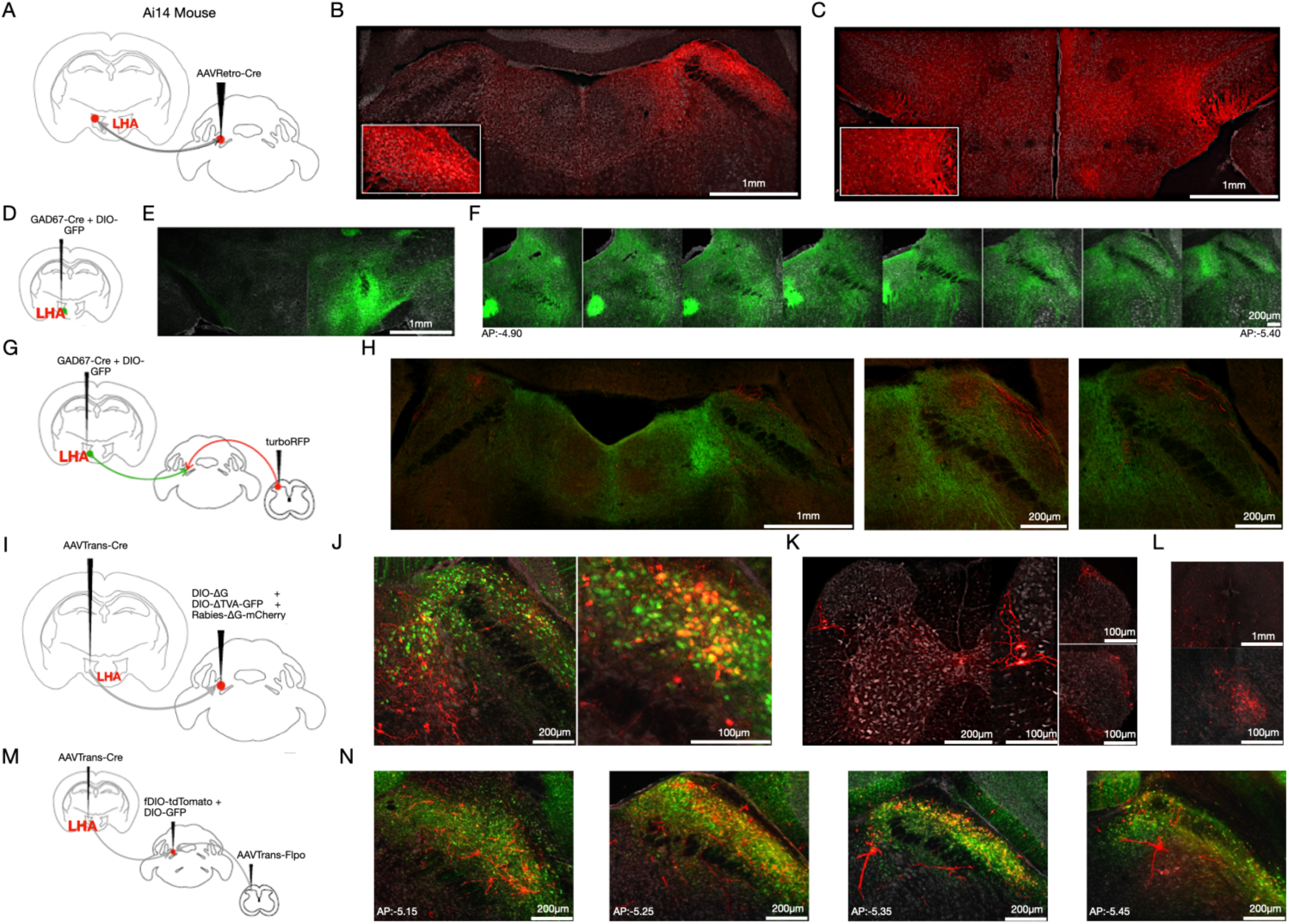
Spinal and lateral hypothalamic inputs synapse on the same LPBN neurons. (A) A schematic of the viral strategy used to identify the inputs LPBN receives using a AAV encoding retrogradely transporting Cre (AAVRetro-Cre) recombinase in Ai14 transgenic mice (n=4). (B) Representative confocal images of the site of AAVRetro-Cre injection in the LPBN showing tdT expression (red). (C) Representative confocal images of the hypothalamic region show all the cells labeled retrogradely from LPBN in the LHA (red). (D) A schematic of the viral strategy used to label the axon projections of the inhibitory lateral hypothalamic (LHA) neurons (n=3). (E) A representative confocal image of the inhibitory LHA cells being labeled with GFP (green). (F) Confocal images of serial LPBN coronal sections arranged in rostral to caudal order (AP:-4.90 to AP:-5.40) getting axon projections from the inhibitory LHA cells. (G) A schematic of the viral strategy used for marking the axon projections LPBN receives from the inhibitory cells in the LHA and the projection neurons in the lumbar region of the spinal cord (n=3). (H) Representative confocal images of the coronal section of the LPBN showing both RFP(red) and GFP (green) projections from the lumbar and LHA respectively. (I) A schematic of the monosynaptic Rabies tracing based strategy used to identify if the cell bodies of LPBN that receive inputs from the LHA are the same ones that also receive input from the lumbar region (n=5). (J) Representative confocal images of the LPBN sections showing the starter cells co-expressing GFP (green) and the rabies cells expressing mCherry (red). (K) Representative confocal images showing the cells in the lumbar region of the spinalcord, especially the superficial laminae neurons labeled with mCherry (red). (L) Representative confocal images showing the cells in the PAG (top) and CeA(bottom) labeled with mCherry (red). (M) A schematic of the viral strategy used to label cells in the LPBN that receive inputs from both the LHA and the lumbar region of the spinal cord. (N) Representative confocal images across the LPBN cell bodies show co-expression of both GFP and tdT (yellow).

Next, we investigated if LPBN neurons that receive LHA inputs also receive synaptic inputs from the spinal cord. Using monosynaptic rabies tracing^18^ (AAV-Trans-Cre in the LHA, AAV-DIO-TVA-GFP, AAV-DIO-ΔG, and Rabies-ΔG-mCherry in the LPBN (Fig. 4J) we labeled LPBN neurons that receive inputs from the LHA (LPBN^LHA^) (green) and neurons that provide monosynaptic inputs to the LPBN^LHA^ neurons (mCherry). Surprisingly, we found mCherry cells in the superficial layers of the dorsal horn of the lumbar spinal cord (Fig. 4K) apart from hind- and mid-brain targets of LPBN in the PAG and central amygdala (Fig. 4L), indicating that LPBN^LHA^ neurons receive direct inputs from the spinal cord. In a complementary intersectional approach, we injected the anterograde trans-synaptically transported recombinases AAV-TransFlpO in the lumbar spinal cord and AAVTrans-Cre in the LHA and recombinase-dependent reporters DIO-GFP and fDIO-tdTomato, in the LPBN. We observed LPBN neurons co-expressing GFP and tdTomato (Fig. 4N), supporting the data from monosynaptic rabies tracing experiments (Fig. 4I) that a subpopulation of LPBN neurons receives inputs from both the LHA and spinal cord. Interestingly, while many GFP-expressing cells did not express tdTomato, most tdTomato cells expressed GFP (Fig. S4), indicating that a greater percentage of LPBN neurons receive input from the LHA than the spinal cord, and most LPBN neurons that receive inputs from the LHA also receive inputs from the spinal cord. Together these data demonstrate that that spinal projection neurons and the descending inhibitory LHA neurons share synaptic targets in the LPBN synapse onto the same cells, which may allow the spinal and hypothalamic inputs simultaneously control the activity of a subpopulation of LPBN neurons.

### 3.5. Stimulation of spinal inputs to the LPBN or the LPBN^spinal^neurons increases coping responses to cold pain

Next, we sought to establish the role of the spinal-PBN pathway in emotional and reflexive coping responses to cold in mice with neuropathic pain. To address this, first, we tested if activation of axon terminals of lumbar spinal neurons in the LPBN alone is sufficient to alter licking and shaking responses to cold in mice with neuropathic pain. To that end, we expressed the light-sensitive Channelrhodopsin2 (ChR2) in the lumbar spinal cord and shined blue light via fiber-optic cannula on the contralateral LPBN to excite the axon terminals (Fig. 5B). Activation of lumbar terminals in the PBN caused an increase in licks but not shakes on the cold-plate test in mice with cold allodynia (Fig. 5C & D), suggestive of an increase in affective-motivational coping responses to noxious cold stimuli. Next, we sought to test if chemogenetic activation or silencing of LPBN neurons that receive synaptic inputs from the lumbar spinal cord (LPBN^spinal^) is sufficient to alter licking and shaking responses to cold in mice with neuropathic pain. To address this, we expressed the excitatory or inhibitory DREADDs, hM3Dq and hM4Di, respectively, in LPBN^spinal^ neurons by injecting AAVTrans-Cre in the spinal cord and DIO-hM3Dq-mCherry (Fig. 5G) or DIO-hM3Dq-mCherry (Fig. 5K) in the LPBN and activated the DREADDs by administering DCZ (i.p). While chemogenetic activation of LPBN^spinal^ neurons increased lick responses to cold in oxaliplatin-injected mice (Fig. 5H & I), chemogenetic silencing of LPBN^spinal^ neurons (Fig. 5L) did not alter licking or shaking (Fig. 5M & N), suggesting that LPBN^spinal^ neurons may play a role in, but are not necessary for expressing emotional responses to cold hypersensitivity. Interestingly, in the cold-puff test, while activating lumbar terminals in the LPBN increased the number of reflexive shakes, engaging or silencing the LPBN^spinal^ neurons did not affect the licks or shakes (Fig. 5E, J & O), demonstrating that activation of spinal terminals in the LPBN and the postsynaptic LPBN^spinal^ neurons promotes licking behaviors in response to sustained but not transient pain stimuli and that activation of LPBN^spinal^ neurons is insufficient to elicit affective-motivational responses in neuropathic mice in response to transient cold stimuli.

**Fig 5. Stimulation.**
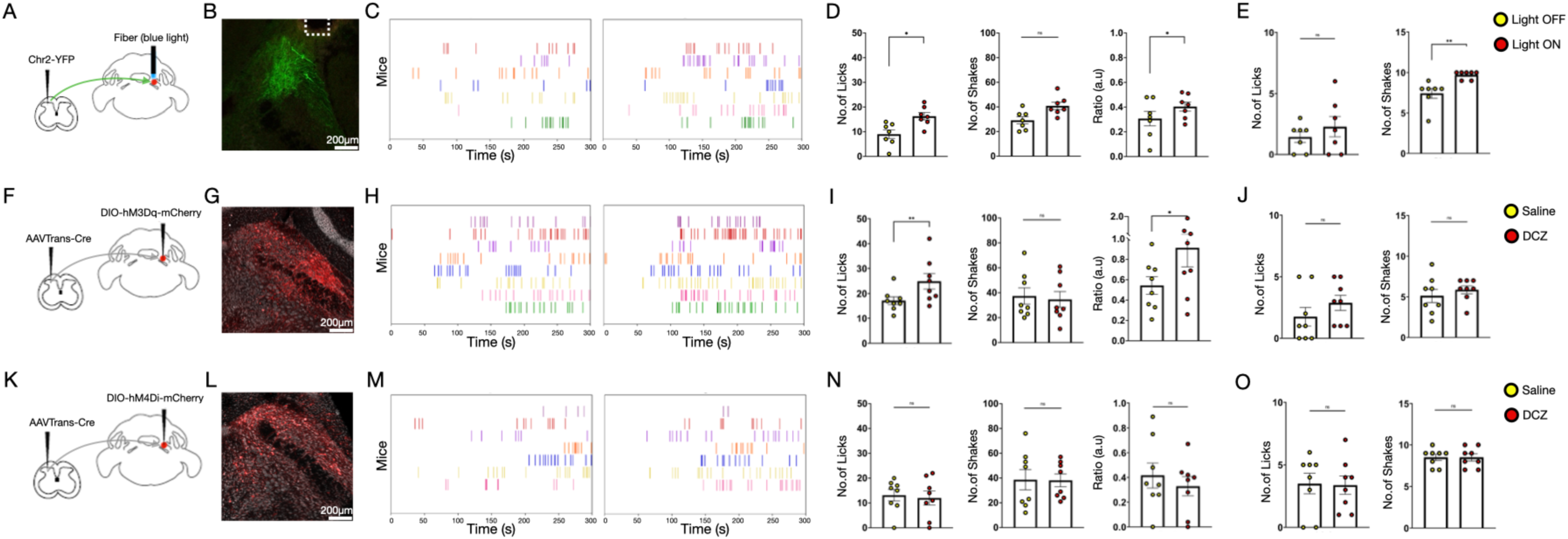
of spinal inputs on LPBN and LPBN^Spinal^ neurons promote licking response. (A) A schematic of the viral strategy used to activate the spinal cord inputs to thecontra LPBN. (n=7). (B) A representative confocal image to show the expression of ChR2-YFP (green) of the spinal cord inputs in the LPBN and fiber location. (C) Raster plots depicting the licks during the cold plate assay where each row is from amouse with light OFF (left) and with light ON (right). (D) Optogenetic activation of lumbar inputs to LPBN during the cold plate assay after i.pl. oxaliplatin, comparing the frequency of licks (left) with light OFF (9.00 ± 1.75) and ON (16.29 ± 1.49) (\**p* = 0.019); the frequency of shakes (middle) with light OFF (29.14 ± 2.91) and ON (40.86 ± 2.92); and the lick-to-shake ratios (right) with light OFF (0.31 ± 0.06) and ON (0.40 ± 0.04) (\**p* = 0.038). (t-test \**p* < 0.05, n=7). (E) Optogenetic activation of lumbar inputs to LPBN after oxaliplatin injection during cold air puff trials, comparing the number of licks (left) with light OFF (1.43 ± 0.43) and ON (2.29 ± 0.84); the number of shakes (right) with light OFF (7.43 ± 0.61) and ON (9.71 ± 0.18) (\*\**p* = 0.0047). (t-test \*\**p* < 0.01, n=7). (F) A schematic of the viral strategy used to activate the lumbar LPBN^Spinal^ neurons (n=8). (G) A representative confocal image to show the expression of hM3Dq-mCherry in the LPBN (red). (H) Raster plots depicting licks during the cold plate assay where each row is from amouse after i.p saline (left) and i.p DCZ (right). (I) Chemogenetic activation of lumbar LPBN^Spinal^ neurons during the cold plate assay after i.pl. oxaliplatin, comparing number of licks (left) after i.p. saline (17.13 ± 1.54) and DCZ (24.88 ± 3.11) (\*\**p* = 0.005); the number of shakes (middle) after i.p. saline (37.38 ± 6.49) and DCZ (34.63 ± 6.32); and the the lick-to-shake ratios (right) after i.p. saline (0.54 ± 0.09) and DCZ (0.91 ± 0.19) (\**p* = 0.012). (t-test \*\**p* < 0.01, and \**p* < 0.05, n=8). (J) Chemogenetic activation of LPBN cells postsynaptic to the lumbar region of the spinal cord during the cold air puff trials, comparing the number of licks (left) after i.p. saline (1.75 ± 0.75) and DCZ (2.88 ± 0.61) and the number of shakes (right) after i.p. saline (5.13 ± 0.81) and DCZ (5.88 ± 0.48) (n=8). (K) A schematic of the viral strategy used to inhibit the lumbar LPBN^Spinal^ neurons (n=8). (L) A representative confocal image to show the expression of hM4Di-mCherry in the LPBN^Spinal^ neurons (red). (M) Raster plots depicting the licks during the cold plate assay where each row is from amouse after i.p saline (left) and after i.p DCZ (right). (N) Chemogenetic inhibition of LPBN^Spinal^ neurons during the cold plate assay after i.pl. oxaliplatin, comparing the number licks (left) after i.p. saline (13.13 ± 2.30) and DCZ (12.00 ± 2.84); the number of shakes (middle) after i.p. saline (38.63 ± 8.17) and DCZ (38.13 ± 5.16); and the number of the lick-to-shake ratios (right) after i.p. saline (0.42 ± 0.10) and DCZ (0.33 ± 0.08) (n=8). (O) Chemogenetic inhibition of LPBN cells postsynaptic to the lumbar region of the spinalcord during the cold air puff trials, comparing the number of licks (left) after i.p. saline (3.50 ± 0.82) and DCZ (3.38 ± 0.73); and the number of shakes (right) after i.p. saline (8.50 ± 0.38) and DCZ (8.50 ± 0.42) (n=8).

### 3.6. Hypothalamic inhibitory neurons restrict the expression of coping responses to cold pain

Having tested the contribution of spinal-PBN pathway, we similarly examined the contribution of the LHA-PBN pathway to nocifensive behaviors (Fig. 6). To address this first we performed targeted activation of LHA inhibitory inputs to the LPBN, by expressing Chr2 under the Gad67 promoter in the LHA followed by stimulation of the terminals in the ipsilateral LPBN with blue light (Fig. 6B). Activation of the LHA terminals in the PBN promoted reflexive shaking responses on the cold plate test (Fig. 6C & D) without altering the licking response, suggesting that when cold pain is persistent, LHA mediated inhibition of the LPBN is sufficient to bias the coping responses towards a reflexive one, but insufficient to suppress the emotional response. In contrast, when the cold stimuli were transient, as in the cold-puff test, licking but not shakes was suppressed when inhibitory inputs from the LHA to the LPBN were activated (Fig. 6E). Thus, when the cold stimuli were withdrawn after the mice with CIPN had reacted, LHA inputs to the LPBN suppressed the emotional coping response. Still, as the stimuli did not persist after the mice had reacted, excessive reflexive coping responses were not recruited to compensate. Surprisingly, when we chemogenetically stimulated the LPBN neurons post-synaptic to the LHA (LPBN^LHA^), the licking behavior was promoted (Fig. 6I), as seen in experiments with LPBN^spinal^ activation (Fig. 5I). In the cold puff test, the LPBN^LHA^ activation did not alter the licking or shaking (Fig. 6J) behaviors. The targeted stimulation of the LPBN^spinal^ and LPBN^LHA^ induces the preferable expression of emotional coping responses over reflexive ones. In addition, we tested if the chemogenetic silencing of the LPBN^LHA^ would alter the nocifensive responses to cold in neuropathic mice and found that both emotional and reflexive coping behaviors remained unaltered (Fig. 6M, N & O).

**Fig 6.**
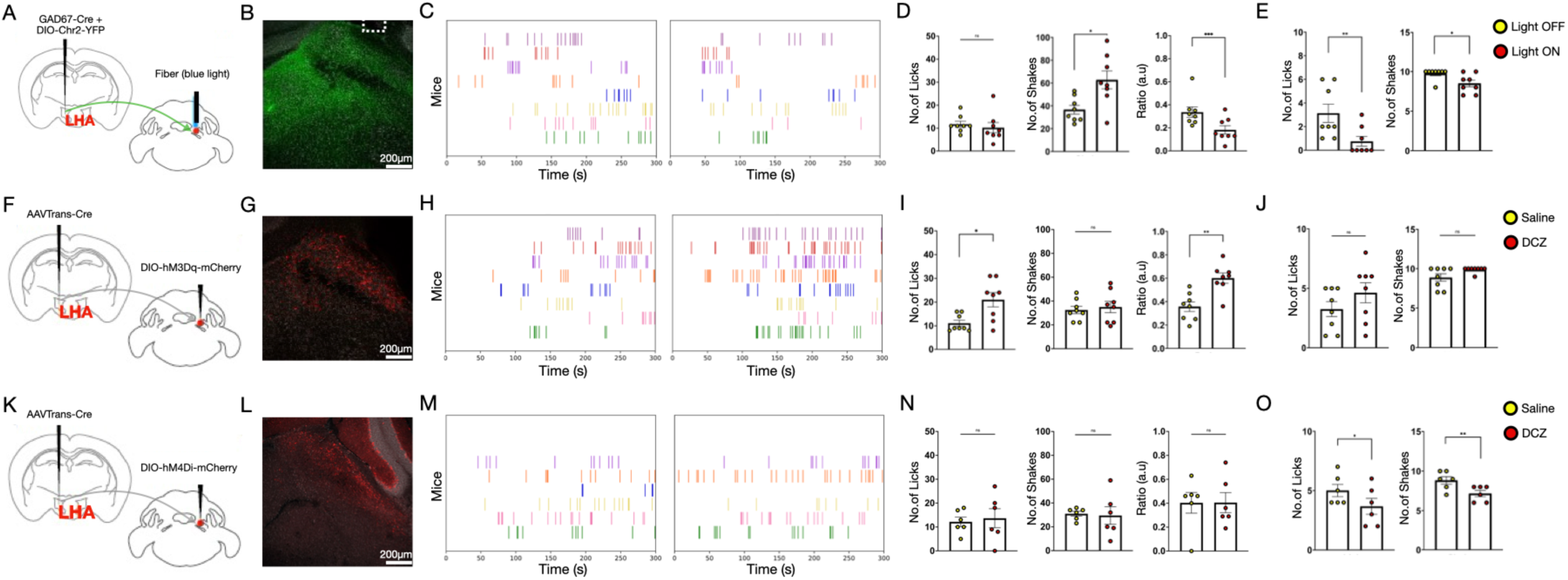
Stimulation of both the inhibitory lateral hypothalamic inputs on LPBN and LPBN^LHA^ promote licking response. (A) A schematic of the viral strategy used to activate only the inhibitory LHA inputs to contra LPBN (n=8). (B) A representative confocal image showing the expression of ChR2-YFP of the inhibitory LHA inputs in the LPBN (green) and the fiber location. (C) Raster plots depicting the licks during the cold plate assay where each row is from amouse with light OFF (left) and light ON (right). (D) Optogenetic activation of inhibitory LHA inputs to the LPBN during the cold plate assay after i.pl. oxaliplatin, comparing the number of licks (left) with light OFF (11.75 ± 1.31) and ON (10.25 ± 2.26); the number of shakes (middle) with light OFF (36.63 ± 3.82) and ON (62.75 ± 7.86) (\**p* = 0.022); and the lick-to-shake ratios (right) with light OFF (0.34 ± 0.047) and ON (0.18 ± 0.04) (\*\*\**p* = 0.0005). (t-test \*\*\**p* < 0.001, and \**p* < 0.05, n=8). (E) Optogenetic activation of inhibitory LHA inputs to the LPBN after i.pl. oxaliplatin during cold air puff trials, comparing the number of licks (left) with light OFF (3.13 ± 0.77) and ON (0.75 ± 0.41) (\*\**p* = 0.002); and the number of shakes (right) with light OFF (8.50 ± 0.42) and ON (9.75 ± 0.25) (\**p* = 0.049). (t-test \*\**p* < 0.01, and \**p* < 0.05, n=8). (F) A schematic of the viral strategy used to activate the LPBN^LHA^ (n=8). (G) A representative confocal image to show the expression of hM3Dq-mCherry in the LPBN (red). (H) Raster plots depicting the licks during the cold plate assay where each row is from amouse after i.p. saline (left) and after i.p DCZ (right). (I) Chemogenetic activation of LPBN cells postsynaptic to the LHA during the cold plate assay after i.pl. oxaliplatin, comparing the number of licks (left) after i.p. saline (11.13 ± 1.26) and DCZ (21.00 ± 3.01) (\**p* = 0.011); the number of shakes (middle) after i.p. saline (32.50 ± 3.11) and DCZ (35.00 ± 4.76); and the lick-to-shake ratio (right) after i.p. saline (0.36 ± 0.04) and DCZ (0.60 ± 0.04) (\*\**p* = 0.0026). (t-test \*\**p* < 0.01, and \**p* < 0.05, n=8). (J) Chemogenetic activation of LPBN^LHA^ during the cold air puff trials, comparing the number of licks (left) after i.p. saline (3.25 ± 0.62) and DCZ (4.63 ± 0.84); and the number of shakes (right) after i.p. saline (8.88 ± 0.48) and DCZ (9.88 ± 0.13) (n=8). (K) A schematic of the viral strategy used to inhibit the LPBN^LHA^ (n=6). (L) A representative confocal image to show the expression of hM4Di-mCherry in the LPBN (red). (M) Raster plots depicting the licks during the cold plate assay where each row is from amouse after i.p saline (left) and after i.p DCZ (right). (N) Chemogenetic inhibition of LPBN^LHA^ during the cold plate assay after i.pl. oxaliplatin, comparing the number of licks (left) after i.p. saline (12.17 ± 1.99) and DCZ (13.67 ± 3.99); the number of shakes (middle) after i.p. saline (31.00 ± 2.03) and DCZ (29.50 ± 7.37); and the lick-to-shake ratios (right) after i.p. saline (0.40 ± 0.09) and DCZ (0.41 ± 0.08) (n=6). (O) Chemogenetic inhibition of LPBN^LHA^ during the cold air puff trials, comparing the number of licks (left) after i.p. saline (5.00 ± 0.52) and DCZ (3.67 ± 0.67) (\**p* = 0.025); and the number of shakes (right) after i.p. saline (8.83 ± 0.48) and DCZ (7.17 ± 0.40) (\*\**p* = 0.0041). (t-test \*\**p* < 0.01, and \**p* < 0.05, n=6).

Next, we asked what the consequence of simultaneous activation of the LHA inhibitory inputs onto the LPBN^LHA^ and LPBN^spinal^ neurons would be (Fig. 7A &B). We previously observed that sustained activation of either LPBN^Spinal^ or LPBN^LHA^ neurons drives licks in response to noxious cold in neuropathic mice (Fig. 5 F-I, 6F-I). While in independent experiments optogenetic stimulation of the LHA inhibitory inputs onto the LPBN promoted the expression of licking behavior compared to shaking (Fig. 6A-D). We found that on the cold-plate test, simultaneous optogenetic activation of LHA terminals and chemogenetic stimulation of the LPBN^spinal^ neurons increased the frequency of licks to cold pain (Fig. 7C, 7E & 7F) as observed in activation of LPBN^Spinal^ or LPBN^LHA^ neurons (Fig. 5 F-I, 6F-I). However, in contrast to when the LHA terminals were stimulated alone (Fig 6A-D), dual activation restricted the switch in behavioral output to cold pain to a reflexive one (Fig. 7C-F). Thus, concluding that the expression of emotional and reflexive coping responses depends on the intrinsic activity of the LPBN neurons, as well as their ascending, and descending inputs.

**Fig 7.**
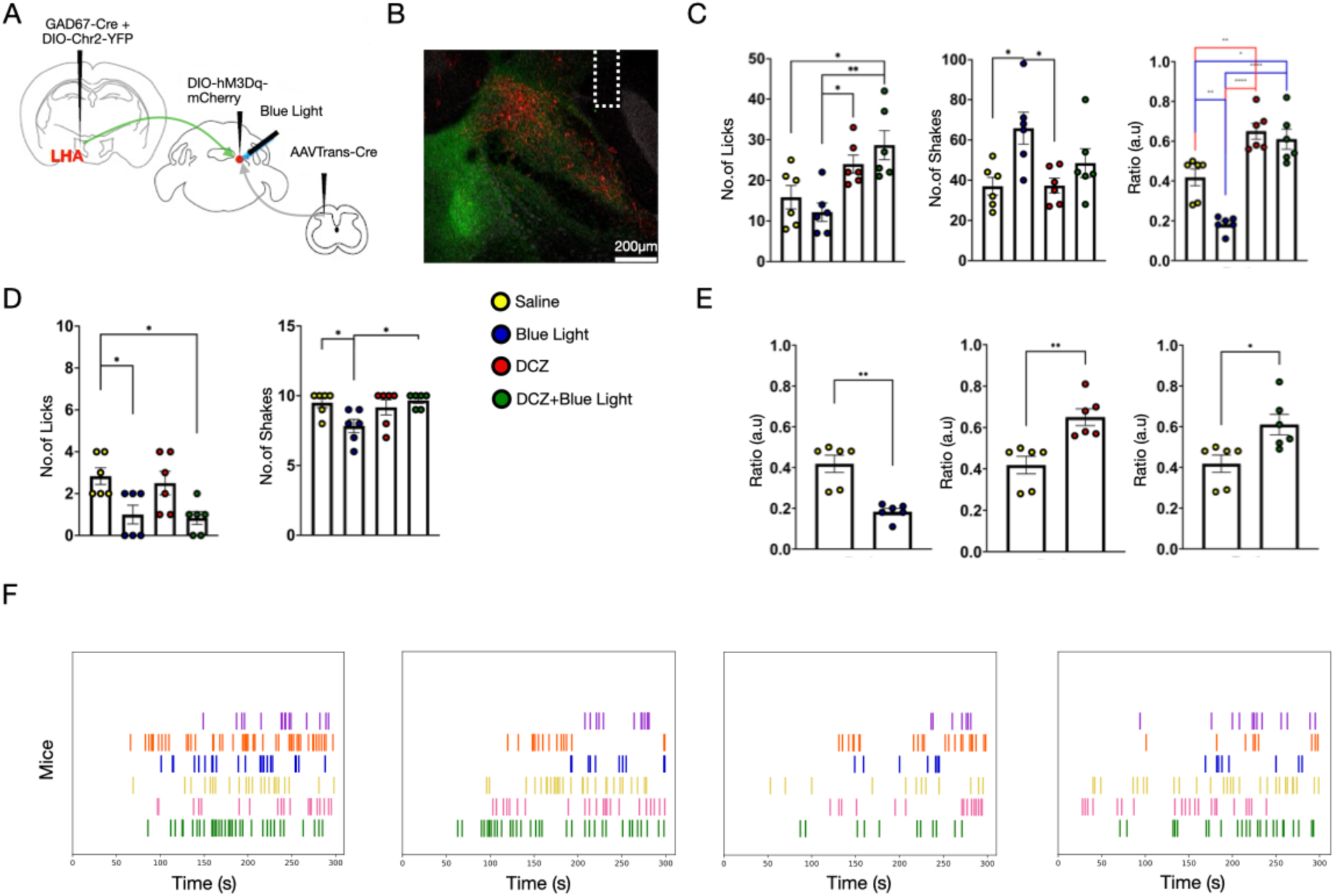
Transient activation of the hypothalamic inputs onto LPBN do not suppress increased licks due to transient LPBN^Spinal^ stimulation. (A) A schematic of the viral strategy used to ontogenetically activate the inhibitory LHA inputs and chemogenetically activate the LPBN cells postsynaptic to the lumbar region of the spinal cord (n=6). (B) A representative confocal image to show the expression of ChR2-YFP of the LHA inputs in the LPBN (green) with the fiber location and expression of hM3Dq-mCherry in the LPBN^Spinal^ neurons (red). (C) During the cold plate assay post-i.pl. oxaliplatin, comparing the change in the number of licks (15.83 ± 2.89) (vs DCZ + Blue light \**p* = 0.019) (left), shakes (37.00 ± 4.37) (vs Blue light \**p* = 0.015) (middle), lick-to-shake ratio (0.42 ± 0.04) (vs Blue light \*\**p* = 0.0022) (vs DCZ \*\**p* = 0.0025) (vs DCZ + Blue light \**p* = 0.0127) (right) during no activation; the number of licks (12.16 ± 2.23) (vs DCZ + Blue light \*\**p* = 0.0024) (left), shakes (65.83 ± 7.96) (middle), lick-to-shake ratio (0.18 ± 0.02) (vs DCZ \*\*\*\**p* < 0.0001) (vs DCZ + Blue light \*\*\*\**p* < 0.0001) (right) during activation of only the inhibitory LHA inputs; licks (24.0 ± 2.21) (vs Blue light \**p* = 0.033) (left), shakes (37.33 ± 3.69) (vs Blue light \**p* = 0.017) (middle), lick-to-shake ratio (0.65 ± 0.04) (right) during activation of only the LPBN^Spinal^ neurons; and licks (28.67 ± 3.58) (left), shakes (48.5 ± 7.15) (middle), lick-to-shake ratio (0.61 ± 0.05) (right) during simultaneous activation of LHA inhibitory inputs onto the LPBN^LHA^ and LPBN^spinal^ neurons. (one way ANOVA *p*\*\*\*\* < 0.0001, *p*\*\* < 0.01 and *p*\* < 0.05, n=6). (D) During the cold air puff assay post i.pl. oxaliplatin, comparing the change in the number of licks (2.83 ± 0.40) (vs DCZ + Blue light \**p* = 0.0206) (vs Blue light \**p* = 0.0365) (left), shakes (9.33 ± 0.67) (right) during no activation; the mean value of licks (1.00 ± 0.45) (left), shakes (9.33 ± 0.67) (vs DCZ + Blue light \**p* = 0.025) (vs Saline \**p* = 0.045) (right) during activation of only the inhibitory LHA inputs; licks (2.50 ± 0.56) (left), shakes (9.50 ± 0.50) (right) during activation of only the LPBN^Spinal^ neurons; and licks (0.83 ±0.31) (left), shakes (9.00 ± 0.82) (right) during simultaneous activation of LHA inhibitory inputs onto the LPBN^LHA^ and LPBN^Spinal^ neurons. (one way ANOVA *p*\* < 0.05, n=6). (E) During the cold plate assay post i.pl. oxaliplatin, the number of the lick-to-shake ratios during activation of only the inhibitory LHA inputs (0.18 ± 0.02) (\*\**p* = 0.0059) (left), activation of LPBN^Spinal^neurons (0.65 ± 0.04) (\*\**p* = 0.0084) (middle) and simultaneous activation of LHA inhibitory inputs onto the LPBN^LHA^ and LPBN^Spinal^ neurons (0.61 ± 0.05) (\**p* = 0.0467) (right) compared with control conditions (0.42 ± 0.04) (t-test \*\**p* < 0.01, and \**p* < 0.05, n=6). (F) Raster plots depicting the licks during the cold plate assay where each row is from a mouse with light ON after i.p DCZ (left); only i.p DCZ (left center); only light ON (right center); and after i.p saline (right).

## 4. Discussion

Here, we sought to understand the circuit underpinnings of behavioral responses to ambient cold in mice with neuropathy-induced cold hypersensitivity. We found that i.pl. oxaliplatin injection caused cold hypersensitivity, promoting reflexive paw shakes and affective-motivational or emotional behavioral responses like licks. We hypothesized that the LPBN neurons might mediate the nocifensive behavioral responses to cold stimuli in mice with peripheral neuropathy. Indeed, we found that in neuropathic mice, the LPBN neurons are sensitized to cold stimuli and, when activated, accentuate shakes and licks to noxious thermal stimuli. Moreover, our data indicate that a subset of LPBN neurons receive nociceptive inputs from the dorsal horn of the spinal cord and are sufficient for increasing lick responses to cold pain. The same neurons operate under the inhibitory influence of lateral hypothalamus (LHA) neurons, and when these inputs are activated, they accentuate the reflexive shake response. In sum, the activity of a subset of LPBN neurons combined with the strength of their inputs play an important role in the expression of coping responses to cold pain.

Platinum-based chemotherapeutic drugs such as oxaliplatin and cisplatin cause progressive peripheral neuropathy and are often a reason for dose limitation or termination of chemotherapy in patients^14,47,11^. Hypersensitivity to cold is one of the distinguishing features of oxaliplatin/cisplatin-induced neuropathy^11,35,48,64^. Recent research has shed light on the peripheral mechanisms of cold hypersensitivity due to neuropathy. However, the central mechanisms remain unclear^47,65,69,88^. Here, we demonstrate that LPBN is vital in expressing defensive behaviors observed in response to cold pain in mice with oxaliplatin-induced neuropathy (Fig.3). In particular, we focused on the licking behavior. Licking injured paws is an affective-motivational behavior and is also interpreted as a coping response^41,56^. Compared to spinal reflex withdrawal or shakes, the brain circuits primarily drive the coping licking behavior. The coping behaviors are translationally relevant, as these behaviors are evoked when the pain is persistent and cannot be resolved by evading or terminating the pain-causing stimuli.

LHA has long been known to mediate pain-induced behavioral responses^23,26,36,52^. LHA-mediated pain modulation is thought primarily through its excitatory projections to the LPBN and PAG^26,77,10,55^. However, the excitatory neurons in the LHA are relatively less in number than the inhibitory ones^45,76^. The role of LHA inhibitory neurons in pain modulation remains poorly understood. Here, we show that the inhibitory LHA inputs to the LPBN can have an effect on the nature of coping response to cold pain in mice. This finding aligns with the known roles of the GABAergic LHA neurons in wakefulness, arousal, robust-consummatory behaviors, reward, and stress^13,16,73,79^, as, these physiological states are known to modulate pain-induced behaviors^3,31,54,63^.

Several recent studies indicate that LPBN neurons act as a gateway for the transmission of nociceptive information to brain regions such as the medulla, central amygdala, hypothalamus, and paraventricular thalamus, driving fear learning, escape response, aversion, and anxiety. However, it has been less clear how the information transmission from the LPBN is gated. Here, we show that the ascending excitatory and descending inhibitory inputs compete at the LPBN and mediate licking response to innocuous cold stimuli in neuropathic mice. Similarly, the frequency of occurrence and the intensity of the behaviors, such as pain-evoked evasion and freezing, can be gated at the LPBN by descending inhibition and ascending activation. Whole-brain monosynaptic input mapping from LPBN^spinal^ neurons can reveal novel inputs that may gate nocifensive behaviors. Interestingly, there are excitatory inputs to the LPBN from LHA and other mid- and fore-brain areas. These inputs along with ascending inputs can accentuate affective-motivational behaviors in animals with chronic pain.

---

**Fig S1.**
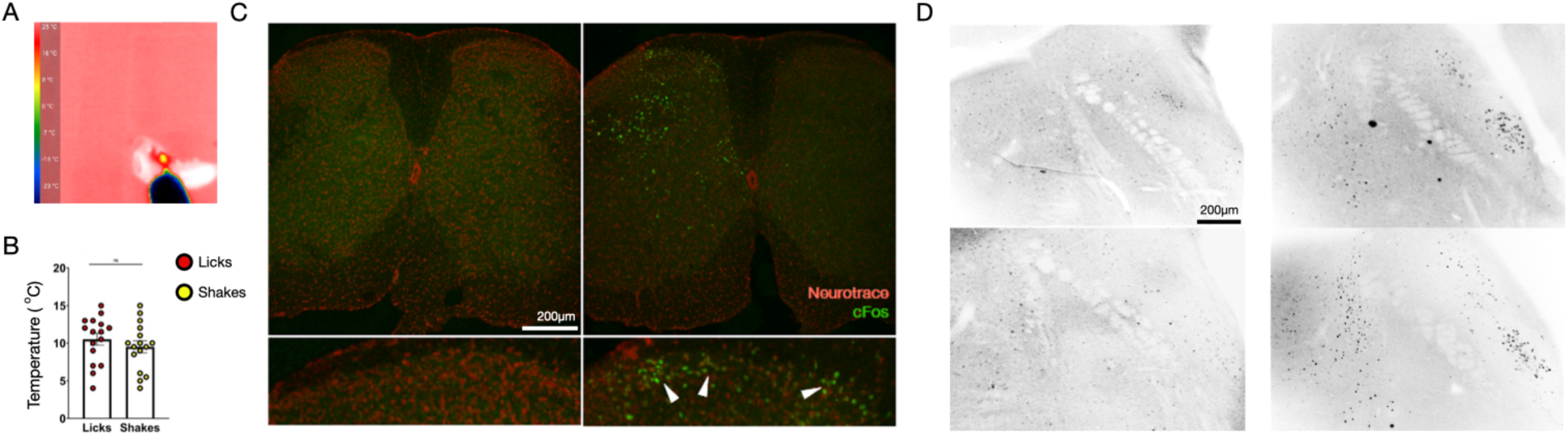
Intraplantar injection of oxaliplatin simulates chronic pain selective to cold temperatures. (A) A representative thermal camera image of a trial during the cold air puff assay. (B) Average temperature at which the mice demonstrated licks (10.5° ± 0.8°) or shakes (9.5° ± 0.8°) post i.pl. oxaliplatin during the cold air puff assay (n=6) were compared. (C) A representative confocal image of the lumbar spinal cord sections from a mouse injected with i.pl. saline (left) and a mouse injected with i.pl. oxaliplatin (right) during a cold plate assay (Neurotrace - red, cFos - green, n=3) (D) Representative confocal images of the LPBN coronal sections from a mouse injected with i.pl. saline (left) and a mouse injected with i.pl. oxaliplatin (right) during a cold plate assay (cFos - Black, n=3).

**Fig S2.**
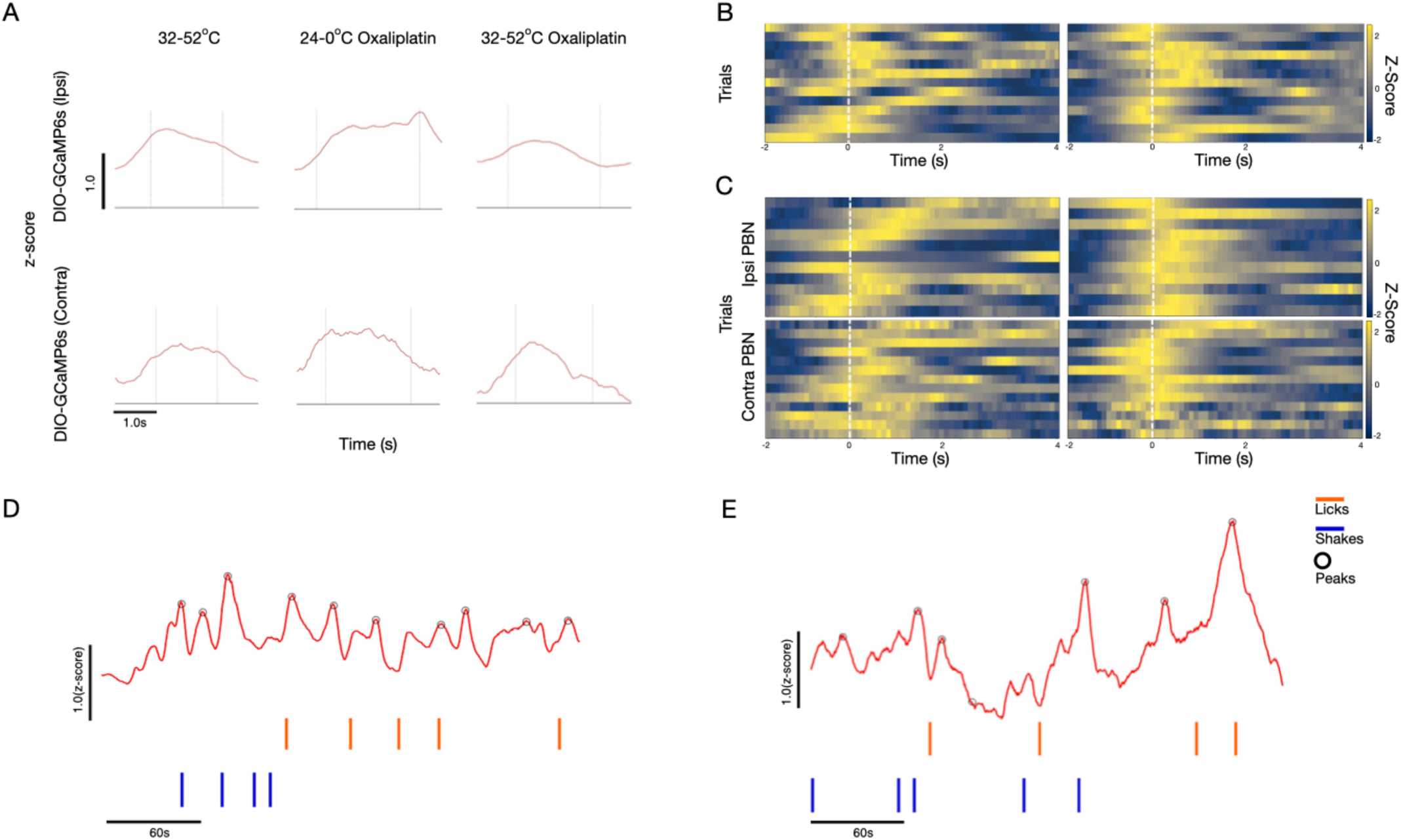
Contralateral and ipsilateral LPBN neurons preferentially respond to cold allodynia. (A) Representative trace graphs of LPBN^VGlut2^ activity during a licking bout which occurs during gradient hotplate assay (left column), cold plate assay after i.pl. oxaliplatin (middle column) and gradient hotplate assay after i.pl. oxaliplatin (right column). The licking bout occurs between the two black lines. The recordings are Z-scores from ipsilateral LPBN^VGlut2^ neurons (top row, n=3) and contralateral LPBN^VGlut2^ neurons (bottom row, n=3). (B) Representative heatmap trials for ipsilateral licks (left) and contralateral licks (right) during gradient hotplate assay with normalized Z-Score values 2 seconds before the start of the responses and 4 seconds after the onset of responses. The white dotted lines indicate the onset of licks. (trials =13, n=3). (C) Representative heatmap trials for licks (left column) and shakes (right column) with normalized Z-Score values 2 seconds before the start of the responses and 4 seconds after the onset of responses. The white dotted lines indicate the start of the responses. The trials were recorded during cold plate assay from contralateral LPBN^VGlut2^ neurons (top row) and ipsilateral LPBN^VGlut2^ neurons (bottom row). (trials =13, n=3). (D) A representative trace of Z-Score recorded from ipsilateral LPBN^VGlut2^ neurons and (E) contralateral LPBN^VGlut2^ neurons during the cold plate assay after i.pl. oxaliplatin. The circles on the trace graph are the identified peaks. The orange and blue lines below are timestamps for licks and shakes respectively.

**Fig S3.**
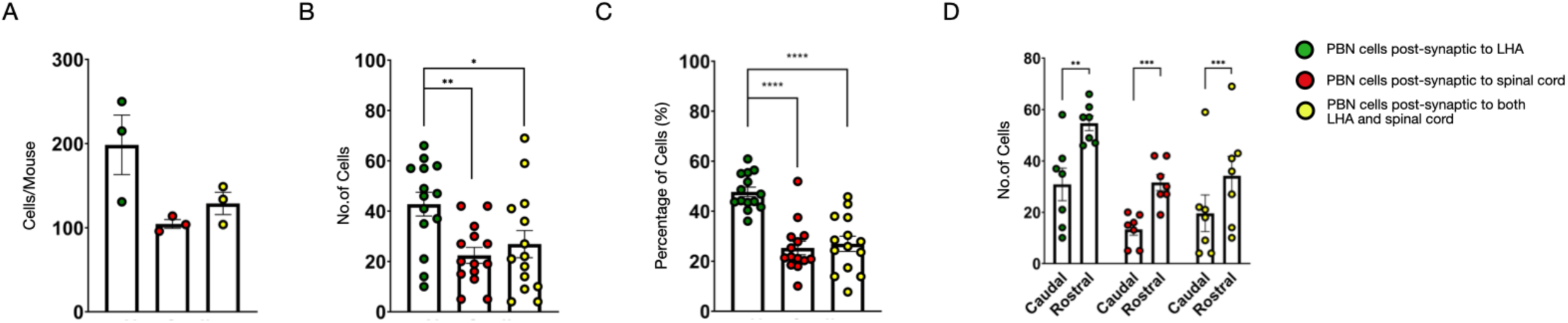
Quantification of LPBN cells receiving inputs from the spinal cord and/or lateral hypothalamus. (A) The total number of LPBN cells getting inputs from the LHA (198.67 ± 35.31), thelumbar region of the spinal cord (104.67 ± 5.21), and from both regions (129.00 ± 13.23) counted across 3 mice. (B) The total number of LPBN cells getting inputs from the LHA (42.79 ± 4.72), the lumbar region of the spinal cord (22.43 ± 3.16) (*p*\*\* = 0.008), and from both regions (26.93 ± 5.40) (*p*\* = 0.045) counted from each coronal section across 3 mice (one way ANOVA *p*\*\* < 0.01 and *p*\* < 0.05, n=3). (C) The total percentage of LPBN cells getting inputs from the LHA (47.73 ± 1.92%) (*p*\*\*\*\* < 0.0001), the lumbar region of the spinal cord (25.31 ± 2.70%) (*p*\*\*\*\* < 0.0001), and from both regions (26.96 ± 3.04%) counted from the labeled cells from each coronal section across 3 mice (one way ANOVA *p*\*\*\*\* < 0.0001, n=3). (D) The total number of caudal (after AP:-5.15) and rostral (before AP:-5.15) LPBN cells getting inputs from the LHA (30.86 ± 6.38 and 54.71 ± 2.89 respectively) (*p*\*\* = 0.0023), the lumbar region of the spinal cord (13.29 ± 2.32 and 31.57 ± 3.18 respectively) (*p*\*\*\* = 0.0001) and from both regions (19.57 ± 7.17 and 34.29 ± 7.53 respectively) (*p*\*\*\* = 0.0006) counted from each coronal section across 3 mice (t-test *p*\*\*\* < 0.001 and *p*\*\*<0.01, n=3).

**Fig S4.**
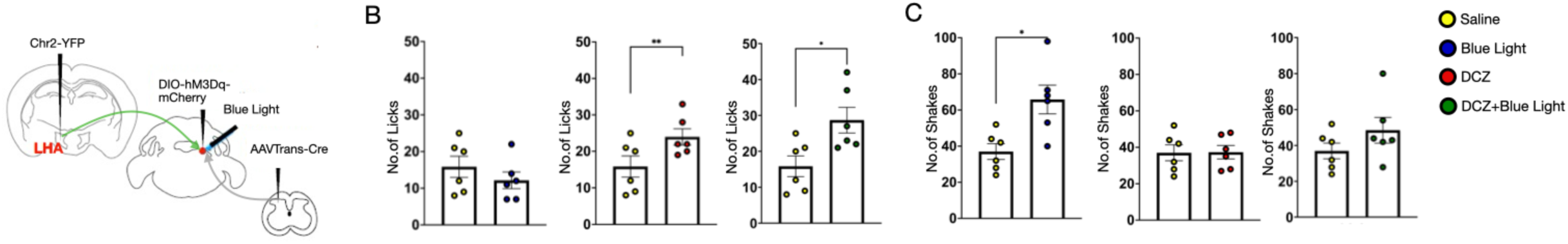
Simultaneous activation of LPBN^Spinal^ neurons and hypothalamic inputs on LPBN resembles the activation from spinal inputs behaviorally on the cold-plate test. (A) A schematic of the viral strategy used to optogenetically activate the inhibitory LHA inputs and chemogenetically activate the lumbar LPBN^Spinal^ neurons (n=6). (B) During the cold plate assay post i.pl. oxaliplatin, the number of the licks during activation of only the inhibitory LHA inputs (12.16 ± 2.23) (left), activation of only the LPBN^Spinal^ neurons (24.0 ± 2.21) (\*\**p* = 0.0067) (middle), simultaneous activation of both populations of cells (28.67 ± 3.58) (right) were compared with control conditions (15.83 ± 2.89) (\**p* = 0.0456) (t-test *p*\*\* < 0.01 and *p*\* < 0.05, n=6).

